# The genetic structure of Norway

**DOI:** 10.1101/2020.03.20.000299

**Authors:** Morten Mattingsdal, S. Sunna Ebenesersdóttir, Kristjan H. S. Moore, Ole A. Andreassen, Thomas F. Hansen, Thomas Werge, Ingrid Kockum, Tomas Olsson, Lars Alfredsson, Agnar Helgason, Kári Stefánsson, Eivind Hovig

**Affiliations:** Centre for Coastal Research, Department of Natural Sciences, University of Agder, Kristiansand, Norway; deCODE Genetics/AMGEN, Inc., Reykjavik Iceland; Department of Anthropology, University of Iceland, Reykjavik, Iceland; NORMENT, Division of Mental Health and Addiction, Oslo University Hospital, Oslo, Norway; Institute of Clinical Medicine, University of Oslo, Oslo, Norway; Institute of Biological Psychiatry, Copenhagen Mental Health Services, Copenhagen, Denmark; Danish Headache Center, Department of Neurology, Copenhagen University hospital, Glostrup, Denmark; Department of Clinical Medicine, University of Copenhagen, Copenhagen, Denmark; The Lundbeck Foundation Initiative for Integrative Psychiatric Research, iPSYCH, Copenhagen, Denmark; Center for Molecular Medicine, Department of Clinical Neuroscience, Neuroimmunology Unit, Karolinska Institutet, Stockholm, Sweden; Institute of Environmental Medicine, Karolinska Institutet, Stockholm, Sweden; Department of Tumor Biology, Institute for Cancer Research, Oslo University Hospital, Oslo, Norway; Faculty of Medicine, University of Iceland, Reykjavik, Iceland; Center for bioinformatics, Department of Informatics, University of Oslo, Oslo, Norway

**Keywords:** population structure, Norway, Scandinavia, Sami, isolation, haplotypes, genetic drift

## Abstract

The aim of the present study was to describe the genetic structure of the Norwegian population using genotypes from 6369 unrelated individuals with detailed information about places of residence. Using standard single marker- and haplotype-based approaches, we report evidence of two regions with distinctive patterns of genetic variation, one in the far northeast, and another in the south of Norway, as indicated by fixation indices, haplotype sharing, homozygosity and effective population size. We detect and quantify a component of Uralic Sami ancestry that is enriched in the North. On a finer scale, we find that rates of migration have been affected by topography like mountain ridges. In the broader Scandinavian context, we detect elevated relatedness between the mid- and northern border areas towards Sweden. The main finding of this study is that despite Norway’s long maritime history and as a former Danish territory, the region closest to mainland Europe in the south appears to have been the most isolated region in Norway, highlighting the open sea as a barrier to gene flow.

## Introduction

Population sub-structures can give rise to false positive associations in association studies of genetic variants (1), can reveal historical patterns of population movements (2, 3), and estimates of ancestry have potential in informing genealogy and forensic genetics (4). Natural features, such as the sea and mountain ridges, tend to limit gene flow between groups of individuals (5), resulting in reproductive isolation and divergence in allele frequencies over time. This divergence may be especially pronounced in smaller populations, due to greater genetic drift. Among the populations in Northern Europe, geographically structured differences are primarily due to isolation by distance, but may also result from founding effects and subsequent isolation (6, 7). Further, isolation and reduction of gene flow within a geographical area can also manifest an increase in recessive Mendelian disorders (8, 9) and founder mutations. Indeed, geographically clustered and expanding BRCA1 founder mutations have been previously reported for Norway (10, 11).

Norway is one of the most sparsely populated countries in Europe, but little is known about its main genetic structure. Its relatively large landmass has the longest coastline in Europe, but has a population of only ~5 million, that includes one of the few indigenous peoples of Europe, the Sami. With unfavorable climatic conditions, combined with the third least arable land in Europe, Norway has provided its people with limited agricultural opportunities. Historically, farms were fragmented through inheritance to ever smaller units, ultimately resulting in unsustainable population growth, especially during the 19^th^ century. Combined with poverty, this motivated the mass emigration of a substantial fraction (1/3) of the population to the Americas during the 19th century, a fraction only surpassed by Ireland (12). Despite recent urbanization, leading to one third of the population residing in cities with >100 000 inhabitants, Norway remains characterized by rural communities and small coastal cities. The diversity in dialects across the country suggests limited gene flow in the past (13).

As might be expected, genetic studies show that contemporary Norwegians are most closely related to the neighboring populations of Sweden and Denmark (14, 15). Genetic studies of the human populations of Denmark, Sweden, Finland and Iceland have revealed some intriguing results, highlighting the impact geography has on human genetic variation and admixture, including minimal structure in the Danish population (15), a north-south gradient in Sweden (16) and founder effects and genetic drift in Finland (6, 17) and Iceland (14, 18, 19).

Here, we describe the geographical structure of the Norwegian gene pool in detail, based on microarray genotypes from 6369 unrelated individuals, who were assigned geographical coordinates based on postal codes. As the mean age of these individuals is approximately 64 years, our analysis provides an overview of stratification in the Norwegian gene pool prior to recent episodes of immigration (20, 21).

## Materials and Methods

### Samples

The dataset was derived from a biobank of approximately 18,000 EDTA-contained blood samples collected over a period of 25 years, as a patient self-referral initiative for overrepresentation of cancer in families, with both clinical and research intent. It includes information about family structure and postcodes, which were converted into longitude and latitude coordinates (22). The biobank consists of families, as well as unrelated individuals, with partial pedigree information covering more than 50,000 individuals (10, 11). Its clinical aim was to provide benefit to patients from the established follow-up examinations aiming at early diagnosis and treatment. All participants provided separate written informed consent to the current research, and the study was approved by the regional ethical review board (REK sør-øst C: 2015/2382).

### Genotypes and sample quality control

DNA was extracted and genotyped at deCODE genetics using the Illumina OmniExpress 24 v 1.1 chip, containing assays for 713,014 SNPs. Data analyses were performed both on the “Services for sensitive data” (TSD) platform at the University of Oslo and at deCODE genetics. The genotyped samples were subjected to quality control and processing in the following order (Supplementary Table S1), using PLINK (v1.90b3) (23). First, autosomal SNPs with a missing rate >2% were removed, followed by removal of SNPs with a minor allele frequency (MAF) <2%. Next, samples with more than 2% missing data were excluded, along with those without a postal code. This resulted in 583,183 autosomal SNPs typed in 14,429 individuals remaining. Finally, we identified all pairwise relationships between individuals using the “--related --degree 3” parameter in KING (v 1.2.3) (24), and discarded individuals related up to the third degree, keeping the oldest individual in each lineage. This resulted in a dataset of 6545 individuals with no close relations (kinship coefficient <0.044) and a mean age of 64 years. There was a predominance of females (81%) as the samples were collected through self-referrals for breast cancer.

As our focus is on population events that occurred prior to the 20^th^ century, we performed analyses to exclude individuals from our sample who derive from recent migration from distant populations. We assessed the extent of European (CEU), Asian (CHB) and African (YRI) ancestry in our Norwegian sample using ADMIXTURE (v 1.3.0) (25). After examining the resulting distributions, we set the maximum threshold for African ancestry to 5%, leading to an exclusion of 65 individuals. The extent of Asian ancestry in our dataset was more pronounced (n=141 > 5% Asian). As many of these samples were found to be from the northernmost county of Finnmark, particularly from the Sami town of Kautokeino, we decided to set the Asian ancestry cutoff threshold > 35% (excluding 29 samples), in order to retain individuals of presumed Sami ancestry. To determine if these indeed were of Sami ancestry, we merged our dataset with a public dataset with genotypes from individuals from a range of countries including one known Sami sample (26) and conducted a PCA. In total, we excluded 94 samples from further analysis that exceeded the thresholds of African (>5%) and East Asian ancestry (>35%).

### Sample density

The samples in this study were distributed over most of Norway, with an over-representation of the south-eastern region that houses half the population, and an underrepresentation from the counties of Sogn og Fjordane and Finnmark (Table 1). For most analyses, we assigned individuals to one of the 19 counties of Norway based on postcodes and applied a restriction of a maximum of 200 random samples per county.

**Table 1:** Summary statistics per county. **N**= the number of samples passing quality control. **N\***=the final number of random samples per county included in the final analysis, with max 200. **Mean ROH**=mean sum of Runs-of-Homozygosity in cM. **Mean IBD**=Mean within-county IBD sharing in cM. **Ne**= estimate of effective population size at g=5 ago. **Pop. size and pop. pr. km^2^**= census population size in 1970.

| County | Abb | N | N* | Median<br>sum of<br>ROH | Mean<br>sum<br>of<br>IBD | Ne | Pop<br>pr. km<sup>2</sup> | Pop | Ne / pop |
| --- | --- | --- | --- | --- | --- | --- | --- | --- | --- |
| Østfold | OF | 388 | 200 | 5.5 | 5.7 | 396,000 | 56 | 221,386 | 1,79 |
| Akershus | AK | 1132 | 200 | 5 | 5.2 | 919,000 | 70 | 324,390 | 2,83 |
| Oslo | OS | 913 | 200 | 4.9 | 4.7 | 579,000 | 1127 | 481,548 | 1,20 |
| Hedmark | HE | 325 | 200 | 8 | 8.4 | 93,600 | 6 | 179,204 | 0,52 |
| Oppland | OP | 294 | 200 | 7.5 | 8.1 | 89,100 | 7 | 172,479 | 0,52 |
| Buskerud | BU | 388 | 200 | 5.6 | 7 | 204,000 | 14 | 198,852 | 1,03 |
| Vestfold | VE | 417 | 200 | 6 | 6.1 | 115,000 | 81 | 175,402 | 0,66 |
| Telemark | TE | 240 | 200 | 6.7 | 9.4 | 91,400 | 11 | 156,778 | 0,58 |
| Aust-Agder | AA | 158 | 152 | 8.2 | 10.2 | 118,000 | 9 | 80,839 | 1,46 |
| Vest-Agder | VA | 252 | 200 | 12 | 13.5 | 44,100 | 18 | 124,171 | 0,36 |
| Rogaland | RO | 225 | 200 | 8.4 | 14.2 | 27,600 | 31 | 268,682 | 0,10 |
| Hordaland | HO | 52 | 52 | 8.1 | 7.1 | 55,500 | 25 | 260,492 | 0,21 |
| Sogn og Fjordane | SF | 22 | 22 | 10.5 | 14.8 | 12,000 | 5 | 100,933 | 0,12 |
| Møre og Romsdal | MR | 187 | 187 | 7.8 | 9.6 | 270,000 | 15 | 223,709 | 1,21 |
| Sør-Trøndelag | ST | 1011 | 200 | 6.7 | 8.7 | 187,000 | 13 | 234,022 | 0,80 |
| Nord-Trøndelag | NT | 187 | 187 | 8.3 | 9.2 | 116,000 | 5 | 117,998 | 0,98 |
| Nordland | NO | 100 | 100 | 6.6 | 8 | 57,400 | 6 | 240,951 | 0,24 |
| Troms | TR | 54 | 54 | 8.8 | 11.5 | 25,600 | 5 | 136,805 | 0,19 |
| Finnmark | FI | 30 | 30 | 27 | 52.2 | 2600 | 2 | 39,757 | 0,07 |
| All |  | 6374 | 2984 | 6.8 | - | - | 12 | 3 888 305 | - |

### Scandinavian dataset

The Norwegian dataset was merged with extended versions of the Danish and a Swedish reference samples used in (14), genotyped on the same genotyping platform. SNPs passing quality control and filtering criteria in the Norwegian dataset were extracted from the Danish and Swedish datasets, expanding the dataset with 1853 Danish and 7966 Swedish samples.

### Principal component analysis

The LD-pruned dataset (PLINK: --indep-pairwise 200 25 0.2”), and specifically excluding 24 high LD regions (27, 28), was subjected to principal component analysis (PCA) as implemented in the eigensoft v6.0.1 (7) function of smartPCA. The pairwise F_ST_ was calculated without automatic removal of outliers (29).

### Shared haplotypes and homozygosity

Missing data in the combined Scandinavian dataset were imputed and phased using beagle v.5 (30). Shared haplotypes, also known as Identity-by-descent (IBD) segments, were detected for autosomal chromosomes using RefineIBD (31), using default settings (minimum length: 1.5 cM, lod > 3 in windows of 40 cM). We increased the minimum size of IBD to 3 cM (31) and summed pairwise IBD sharing between all possible pairs of individuals. Pairwise county level ancestry was determined as the mean of the sum of IBD sharing between individuals residing in the counties in question. County information was available for Norway and Sweden, while Denmark was treated as one geographical unit.

The length of homozygous segments (cM) in each individual were summed to provide a measure of genomic inbreeding, the distribution of which was assessed by county (maximum N samples per county = 200, total N = 2984). To create a smoothed contour map of Norway, we combined the sum of homozygous content per individual with latitude and longitude in spatial regression as with in the Krig function in the R package “fields” (2, 32).

### Historical effective population sizes

Temporal changes in effective population sizes can be estimated by the length and distributions of shared haplotypes (IBD) (33). The effective size (*N*_e_) of a population can be assessed from the pattern of genetic variability in its gene pool and is affected by rates of migration and growth (34, 35). Here, we implemented IBDne (33), for each county using IBD segments called by the RefineIBD algorithm (30, 36), assuming a generation time of 30 years (37).

### Estimation of migration rates and directed gene flow

Effective migration rates in Norway were estimated using EEMS (38), using the LD-pruned dataset. A spatial outline of Norway was constructed by representing it as a concave hull using the R package “concaveman”, and the resulting polygon was used as a border descriptor. A dissimilarity matrix using the bundled script “bed2diff” was constructed. The algorithm assigns individuals to the nearest deme, and by using a stepping-stone model, migration rates are estimated between demes. We performed several iterations with 500 demes. As recommended, we adjusted migration, diversity and degrees-of-freedom parameters for a 10-40% acceptance rate. We set the number of burn-in iterations to 500 000 to ensure that the MCMC algorithm converged.

## Results

### Population structure in Norway

We performed a PCA to detect fine-scale population structure using LD-filtered SNPs (n=102,023) (Supplementary Table S1). First, we color-coded the samples in the PCA (Figure 1). The first component (PC1) seems to capture Uralic-associated admixture (Supplementary Figure S1), and variation in the second component (PC2) reflects drift in southern Norway. The geographical distribution of Uralic associated ancestry was quantified for each county using the results from Admixture (Supplementary Figure S2). Potential sources of Uralic ancestry include the indigenous Sami and later immigrating Finnish minorities. We also found evidence that the third (PC3) component captures meaningful geographical information (Figure 1a & 1b). The mean PC1-10 per municipality are supplied in supplementary materials. We assessed the relationships between PCs and geography (latitude and longitude) using a Pearson’s product moment correlation coefficient test. PC1 showed significant (p < 2e-16) correlations with latitude (r = 0.42) and latitude (r = 0.44), as did PC2 (p < 2e-16; latitude r = −0.32, longitude r = −0.16). To further examine the correlation with geography, we color-coded the samples based on county, and inspected the sample distribution in a PCA plot (Figure 1a & 1b). The five postcodes with the largest mean eigenvalues in PC1 (N individuals >1) were: Kautokeino, Nesseby, Nordreisa, Røyrvik and Alta in the northeast and Hægebostad, Hå, Eigersund, Birkenes and Seljord in the South. A table of municipality with mean PC1-10 values is available (https://doi.org/10.6084/m9.figshare.11235803.v1).

**Figure 1:**
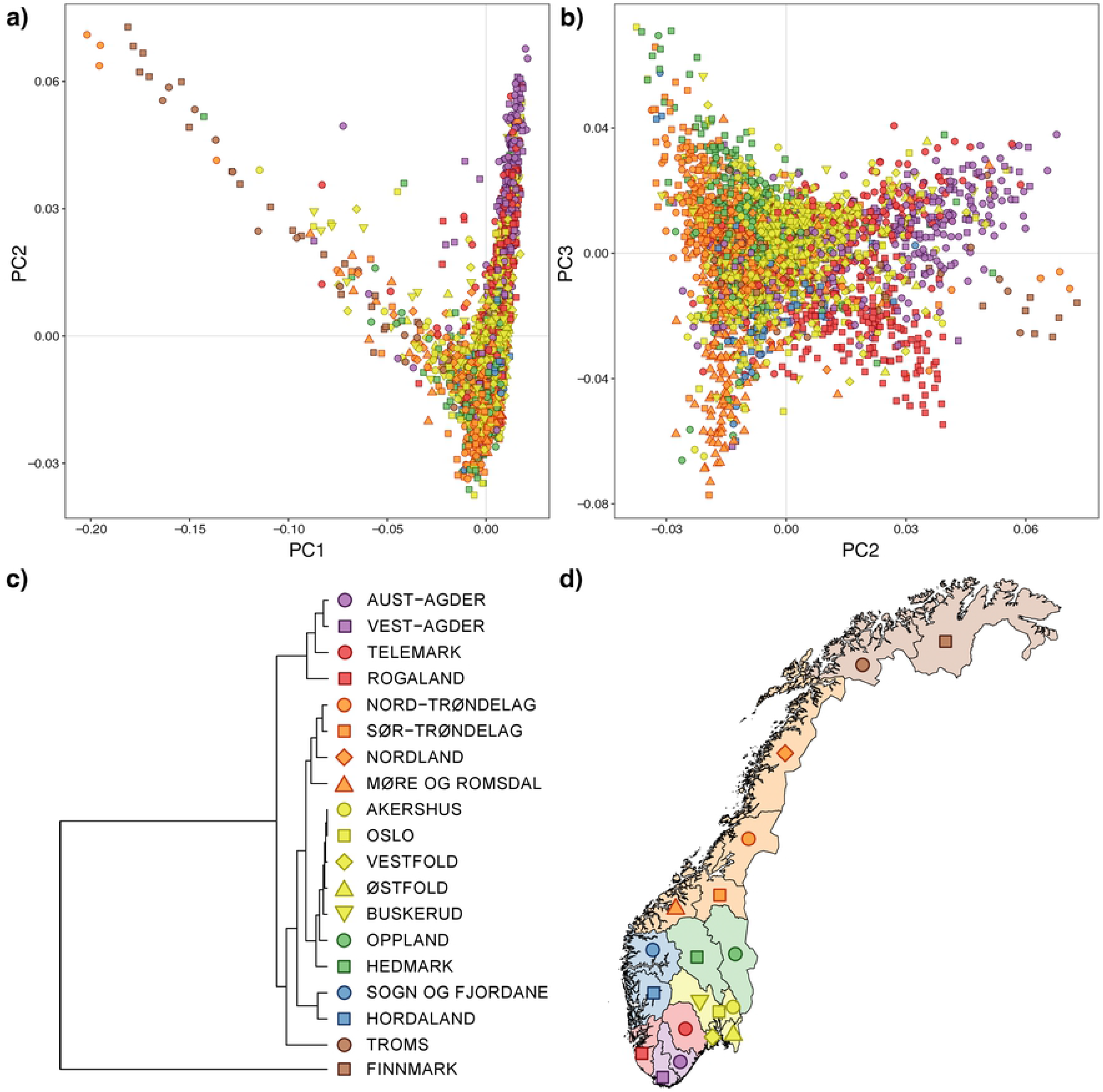
a) & b) PCA plots of LD pruned SNPs (102,023) color-coded by county. The median PC1 and PC2 pr. county is marked with a larger filled circle. PC1 captures the Sami component, and PC2 a southern component of distinctive drift. c) Color-coded map of the counties in Norway. d) Hierarchical clustering of Reich’s F_st_ values, using squared dissimilarities (ward.D2) presented as a phylogram.

To put the Norwegian population in a Scandinavian context, we conducted a PCA of the combined Scandinavian dataset. Here, the divergence of South Norway is apparent (Supplementary Figure S3). In the first two PCs, there are three dimensions of divergence: Uralic-related ancestry, the Norwegian south, and the Swedish north.

### Genetic distances between Norwegian counties

A hierarchical clustering of pairwise F_ST_ distances between counties revealed a similar pattern as the PCA, with the largest divergence in Finnmark in the north, followed by the southern counties of Rogaland, Agder and Telemark (Figure 1b). We note that the counties Møre og Romsdal, Trøndelag and Nordland group together, and that the counties by the Oslofjord area also form a cluster. The average pairwise F_ST_ between Norwegian counties was 0.0012 (max: 0.0073). For comparison, the mean pairwise F_ST_ values for regional differentiation in surrounding countries are: 0.0024 in Finland (max: 0.006), 0.0002 in Denmark, 0.0012 in Sweden (max: 0.0025) and 0.0007 in Great Britain (max: 0.003) (3, 15–17) (all F_ST_ values are derived from the same software, except for the Danish study). Clearly, Finland stands out in this context, and Norway is comparable with Sweden in terms of inter-county differentiation. However, Norway has the largest extent of differentiation within a nation, with Rogaland vs Finnmark, F_ST_ = 0.0073, which is also the most spatially distant (~1250 km) pairwise comparison in Scandinavia (we note that the Swedish study excluded samples with Uralic related ancestry) (16).

### Kinship and inbreeding in Norwegian counties

We assessed the mean autosomal haplotype sharing (IBD > 3cM) within and between counties (Figure 2). By far the greatest within-county mean haplotype sharing was observed in Finnmark (52.2 cM), followed by Sogn og Fjordane (14.8 cM), Rogaland (14.2 cM), and Vest-Agder (13.5 cM). The marked haplotype sharing in Finnmark stands out in a Norwegian context, but elevated haplotype sharing has also been found in the Finnish population, especially eastern Finland (~45 cM) (39), suggesting homogeneity and small effective population sizes. Conversely, the smallest within-county haplotype sharing was observed for the capital area of Oslo (4.7 cM), Akershus (5.2 cM) and Østfold (5.7 cM). The greatest haplotype sharing between counties were observed for Troms and Finnmark in the North (18 cM), and for Vest-Agder and Aust-Agder in the South (10.8 cM).

**Figure 2:**
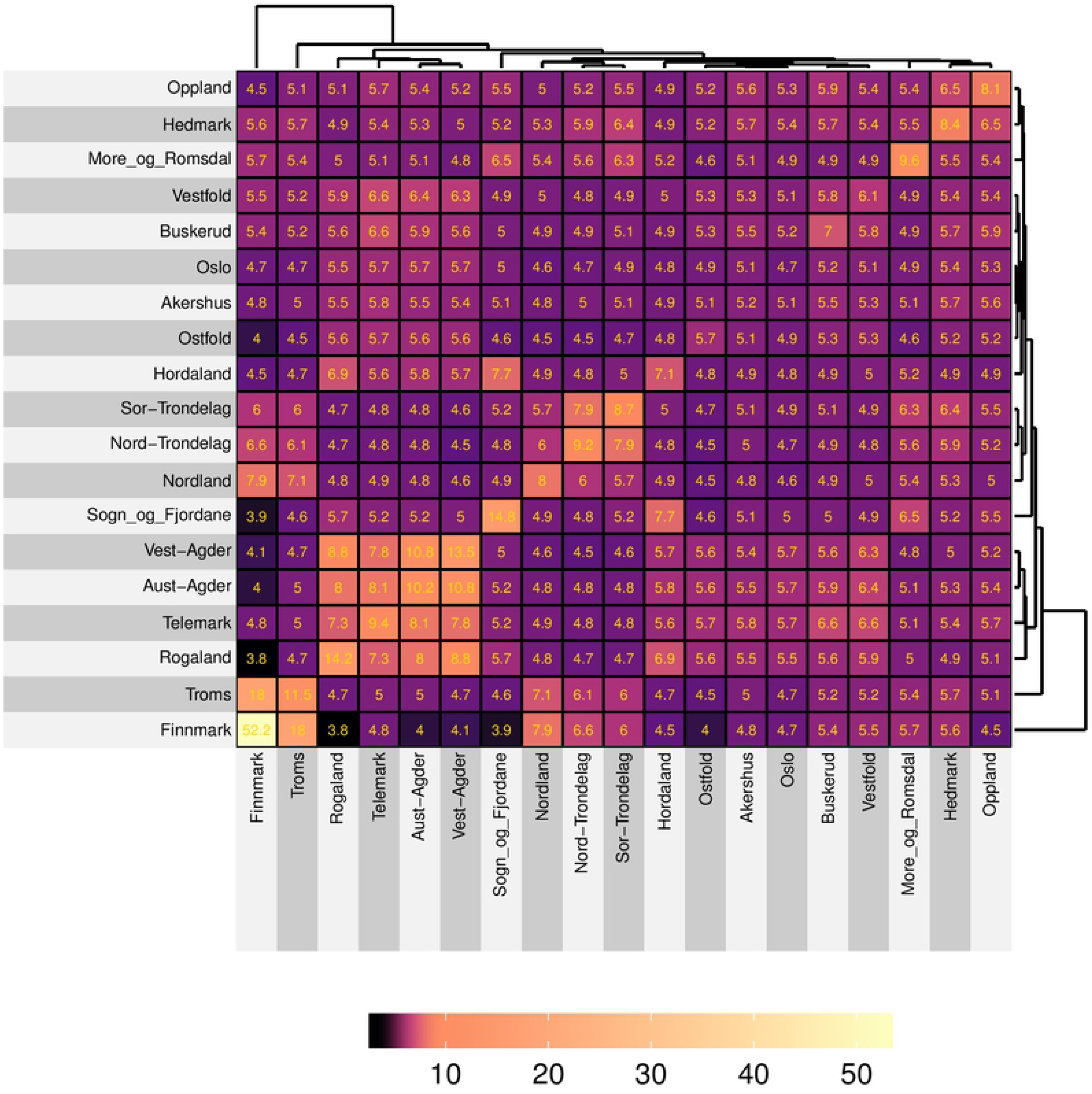
Visual representation and hierarchical clustering of the mean cumulative sum of haplotype sharing (IBD > 3cM) within and between counties in Norway, in centiMorgans (cM). Overall, there is an increased relatedness within the counties (diagonal), and pronounced relatedness between counties form squares.

Homozygosity, measured as the summed length of homozygous segments detected by RefinedIBD, is relatively high in the north, presumably due to increased Sami and Finnish ancestry. Increased homozygosity is also evident in the border areas towards Sweden in the middle, and inland areas of mid-Norway, protruding down to the southwestern coast (Figure 3). Areas with substantially lower degrees of homozygosity include the Oslofjord area in the southeast, the Trondheimsfjord area in the middle, and the northern county of Nordland. The county of Nordland, with no major cities and home to large fishing grounds, appear heterogenous. We also assessed if individuals from rural areas (n=1701) were significantly more homozygous than those from urban areas (20 largest cities, n=1283). Individuals from rural areas were significantly more homozygous than individuals from urban areas, with a median of 6.1 cM and 5.1 cM respectively (two-sided t-test p=9.28×10^−9^).

**Figure 3:**
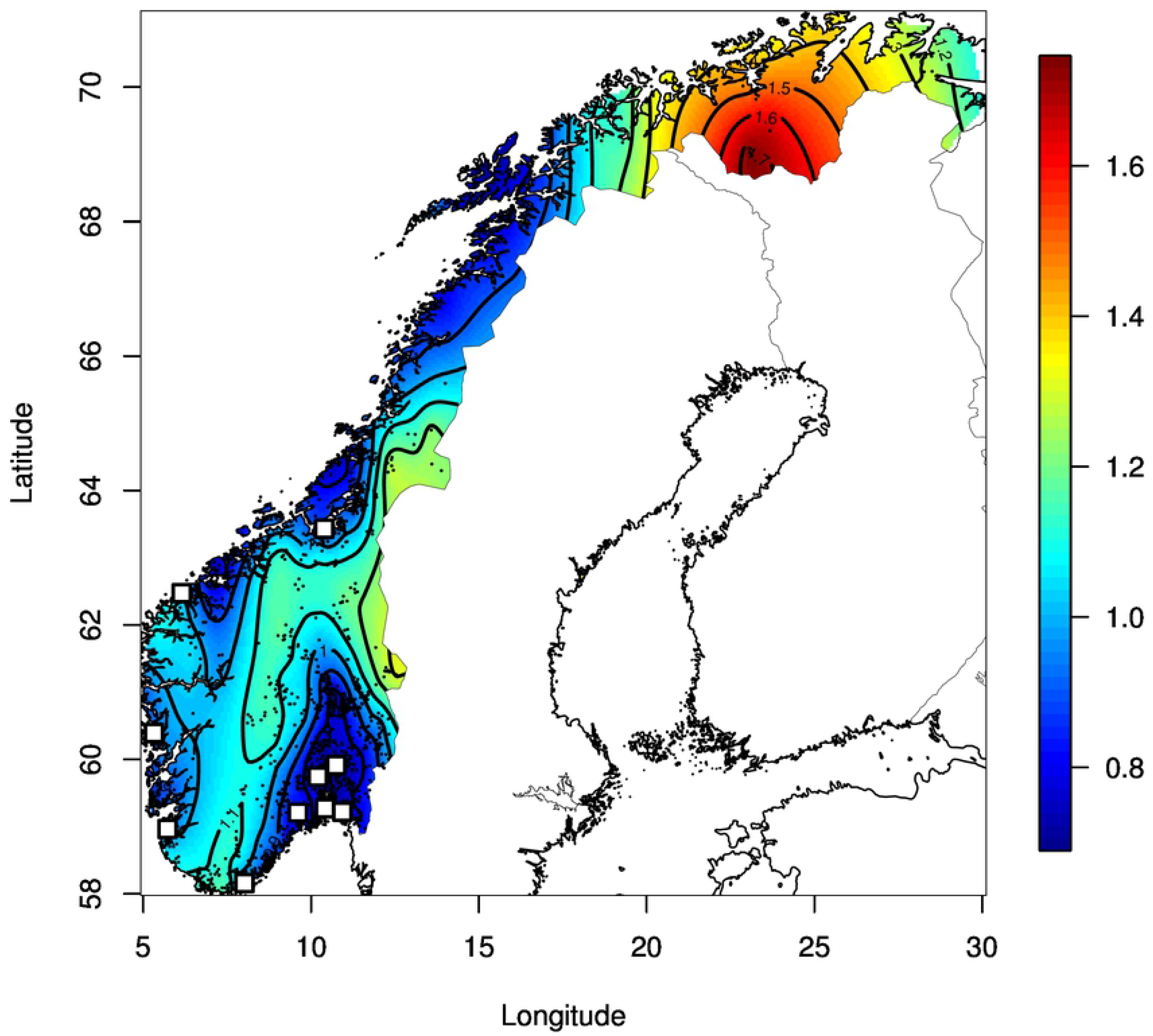
Contour plot of the cumulative sum of homozygous segments (cM) on the log10 scale detected by Beagle, extrapolated by spatial regression (Krig/fields). The black dots represent jittered coordinates of zip codes, using 2984 individuals (max 200 pr. county). The ten most populous cities (> 50 000 inhabitants) are marked with white squares. A continuous belt of elevated homozygosity stretches along in the interior, towards the southwestern coast.

### Kinship to Denmark and Sweden

We explored the mean sum of autosomal haplotype sharing (IBD > 3cM) between Norwegian and Swedish counties, and Denmark as a whole (Supplementary Figure S6 and S7). We find a distinct pattern of low degree of shared ancestry between Norway and Denmark (3.1 cM), including the South/Southeast of Sweden (Skåne=3.3 cM). At the opposite end, the northernmost county in Sweden, Norrbotten, shared 13.1 and 8.1 with Finnmark and Troms, respectively. Further, we detected elevated haplotype sharing between the counties on the border of Norway and Sweden. Noteworthy, the former disputed county of Jämtland, conquered by Sweden in 1679, stands out for having a relatively high IBD sharing with Nord-Trøndelag of 6.6 cM.

### Historical effective population sizes

The distribution of shared IBD segment lengths is also informative about *N*_e_ through time (33, 40). Most, but not all, counties reveal a decrease in effective population sizes, with a minimum around 12-14 generations ago at 1550-1600 AD, assuming a 30-year generation time (Supplementary Figure S4). This minimum has also been reported in other isolated populations in Northern Europe (41).

### Estimation of migrations rates

The simulations of effective migration surfaces returned numerous patterns, some of which were consistent across multiple iterations. These included a general trend of coastal pockets receiving migration and inland barriers (Supplementary Figure S5). We observed three of the notable features. First, was an increased migration rate over a highland area entitled “Hardangervidda” that lies between the two largest cities in Norway, Oslo and Bergen. This genetic corridor corresponds to known ancient trade trails and horse tracks across this highland. Second, there is evidence for barriers in the south, in line with the north-south facing valleys, coinciding with current county borders. Third, we note an isolation of the traditional Sami area of “Finnmarksvidda” in the far north.

## Discussion

We describe for the first time, using common variants, the genetic structure of the Norwegian population at a genome-wide scale. The Sami people, and later immigrating minorities from Finland, like the “Kven” and “Skogfinner” (~1500 AD), are recognized ethnic minorities, and their influence on the genetic landscape of Norway is clearly detectable in the PCA, especially in the three northernmost counties (Figure 1 and Supplementary Figure S1). This is consistent with evidence from a health survey conducted in the 1980’s in Finnmark, where ~25% of the participants reported a Finnish family background. To fully appreciate the extent of Finnish and Sami ancestry, we quantified the extent of Asian ancestry per county (Supplementary Figure S1 & S2). We find a substantial extent of Asian ancestry (mean ~25%, Kautokeino), a size similar to that reported (42) in a single Sami sample (~25% Nganasan). To our knowledge, previous studies of the Sami in Finland report less Asian ancestry (~6%) (43), suggesting a more isolated Sami population in Norway.

Our results further support the divergence, isolation and homogeneity in the southern counties of Norway (Rogaland, Agder and Telemark). The isolation is exemplified by the observation that Oslo has a similar historical profile of effective population size to that of the general British population, while Rogaland had a similar historical profile to the Orkney Islands (41). Further, the counties of Rogaland and Vest-Agder display elevated levels of within-haplotype sharing (~13-14 cM), suggesting isolation and inbreeding (Figure 2), as well as increased homozygosity (Figure 3) and small *N*_e_ (Table 1). This is in line with previous reports on genetic drift in southern Norway (10, 11).

Norway has close historical ties to Denmark, as Norway became a vassal state of Denmark in 1380, lasting 443 years, until 1814. The PCA (Supplementary Figure S4) and IBD analyses strongly suggest that the counties in southern Norway have diverged from the rest of the Norwegian population due to isolation, rather than gene flow from Denmark or some other neighboring populations. We speculate that the isolation in the Norwegian south may be a consequence of an unusual coastline, with an absence of deep fjords, common elsewhere in Norway, the late development of infrastructure like railroad and roads the last 100 years and the region failing to recruit economic migrants.

In a medical context, there is a need to establish national frequency-based databases for disease studies (44). We have taken the first step in this endeavor by documenting geographical patterns of genetic variation in the Norwegian population. Such a database should contain a relatively large amount of frequency differences (weighted F_ST_=0.0073) between geographical regions (Rogaland (200) vs Finnmark (30), weighted F_ST_=0.0073, maximum local F_ST_ = 0.47, rs904274) within Norway. To avoid the undesirable effects of population stratification on genotype-phenotype association studies, and to increase precision, detailed geographical information of individual origin should be included.

For the first time we document restricted gene flow in the southern part of Norway, which is contradicting a commonly held notion of Danish admixture. We next aim to characterize the detailed population structures in the Norwegian population further using rare variants, as rare variants are more geographically clustered, due to their more recent origin.

## Supplemental Data

Supplemental Data include seven figures and one table.

## Acknowledgements

We wish to express our deepest gratitude and respect to the volunteer participants. We also wish to acknowledge Erik Bolstad and ~600 Norwegian volunteers at the “dugnad” at yr.no for collecting and publishing postcodes with coordinates. We also wish to thank Arne Solli for interesting discussions.

## Author Contributions

The study was conceived by E.H, O.A.A, P.M, K.S and A.H. TW and T.F.H. collected the Danish sample and I.K., T.O. and L.A. collected the Swedish sample. Genotyping was performed by K.S. and A.H. Data analysis was performed by M.M., E.H., S.S.E., A.H. and K.H.S.M. The manuscript was drafted by M.M, with contributions from E.H and A.H. All authors commented upon the draft and approved the final manuscript.

## Supporting Information Legends

**Figure S1:** PCA of the dataset from this study (black) merged (SNPs = 58,457) with public datasets (26) of selected and colored European samples, including one single Sami sample (left legend). The size of the black circles (right legend) represents the percentage of Asian ancestry (CHB+JPT) calculated by ADMIXTURE (25).

**Figure S2:** The fraction of Asian ancestry pr. county (mean with standard error of the mean) indicate increased Asian ancestry in the northmost counties of Troms and Finnmark (ADMIXTURE/ HapMap CHB+JPT).

**Figure S3:** PCA plot of 8110 Scandinavian samples, consisting of 2985 Norwegians, 3519 Swedes and 1606 Danes, with regional information. A maximum of 200 samples was set pr. region, and LD pruned (“indep-pairwise 200 25 0.5”), leaving 238,689 SNPs. In additional to the diverging Sami/Finnish samples, samples from the northern counties of Sweden (Norrbotten and Västerbostten) and the southern counties of Norway (Rogaland, Vest-Agder, Aust-Agder and Telemark) display distinctive drift.

**Figure S4:** Changes in effective population sizes though time as estimated by IBDne, using IBD segments > 3 cM and maximum 50 generations back. The upper and lower 95% confidence intervals are marked with dotted lines. Most counties show a decrease in effective population sizes with a minimum around 12-14 generations ago. We assume the decline has been initiated by The Black Plague, with subsequent isolation, having a minimum at 1550-1600 AD (assuming a 30-year generation time). Counties in the far north and far south have the least growth in more recent times.

**Figure S5:** Simulation of effective migration rates using LD-pruned SNPs from 2984 (max 200 pr. county) individuals and 500 demes. Brown indicate areas of significantly reduced migration rates, and blue indicates significantly increased migration on a logarithmic scale. The black circles represent sample size and overlay grid (38).

**Figure S6:** The proportion of shared genomic content between counties in Norway, Sweden and Denmark. The border areas between Norway and Sweden share overall more genetic content compared that of Denmark and southern Sweden.

**Figure S7:** Visual representation and hierarchical clustering of the mean cumulative sum of haplotype sharing (IBD > 3cM) between counties in Norway and Sweden, including Denmark, in centiMorgans (cM). The color-coding does not scale linearly. Overall, Denmark and South/southeastern Sweden share less kinship towards Norway (dark left), than do the bordering counties between Norway and Sweden (upper right).

**Table S1**: Overview of quality control and the retained number of samples and SNPs.

## References

1 Mathieson I, McVean G. Differential confounding of rare and common variants in spatially structured populations. Nat Genet. 2012;44(3):243–6.

2 Karakachoff M, Duforet-Frebourg N, Simonet F, Le Scouarnec S, Pellen N, Lecointe S, et al. Fine-scale human genetic structure in Western France. Eur J Hum Genet. 2015;23(6):831–6.

3 Leslie S, Winney B, Hellenthal G, Davison D, Boumertit A, Day T, et al. The fine-scale genetic structure of the British population. Nature. 2015;519(7543):309–14.

4 Kayser M, de Knijff P. Improving human forensics through advances in genetics, genomics and molecular biology. Nat Rev Genet. 2011;12(3):179–92.

5 Esko T, Mezzavilla M, Nelis M, Borel C, Debniak T, Jakkula E, et al. Genetic characterization of northeastern Italian population isolates in the context of broader European genetic diversity. Eur J Hum Genet. 2013;21(6):659–65.

6 Salmela E, Lappalainen T, Fransson I, Andersen PM, Dahlman-Wright K, Fiebig A, et al. Genome-wide analysis of single nucleotide polymorphisms uncovers population structure in Northern Europe. PLoS One. 2008;3(10):e3519.

7 Patterson N, Price AL, Reich D. Population structure and eigenanalysis. PLoS Genet. 2006;2(12):e190.

8 Abdellaoui A, Hottenga JJ, de Knijff P, Nivard MG, Xiao X, Scheet P, et al. Population structure, migration, and diversifying selection in the Netherlands. Eur J Hum Genet. 2013;21(11):1277–85.

9 Ben Halim N, Ben Alaya Bouafif N, Romdhane L, Kefi Ben Atig R, Chouchane I, Bouyacoub Y, et al. Consanguinity, endogamy, and genetic disorders in Tunisia. J Community Genet. 2013;4(2):273–84.

10 Moller P, Heimdal K, Apold J, Fredriksen A, Borg A, Hovig E, et al. Genetic epidemiology of BRCA1 mutations in Norway. Eur J Cancer. 2001;37(18):2428–34.

11 Moller P, Hagen AI, Apold J, Maehle L, Clark N, Fiane B, et al. Genetic epidemiology of BRCA mutations--family history detects less than 50% of the mutation carriers. Eur J Cancer. 2007;43(11):1713–7.

12 Grytten O. The Economic History of Norway. EH.Net Encyclopedia: EH.Net Encyclopedia; 2008 [Available from: http://eh.net/encyclopedia/the-economic-history-of-norway/.

13 Røyneland U. Dialects in Norway: catching up with the rest of Europe? International Journal of the Sociology of Language2009. p. 7.

14 Ebenesersdottir SS, Sandoval-Velasco M, Gunnarsdottir ED, Jagadeesan A, Guethmundsdottir VB, Thordardottir EL, et al. Ancient genomes from Iceland reveal the making of a human population. Science. 2018;360(6392):1028–32.

15 Athanasiadis G, Cheng JY, Vilhjalmsson BJ, Jorgensen FG, Als TD, Le Hellard S, et al. Nationwide Genomic Study in Denmark Reveals Remarkable Population Homogeneity. Genetics. 2016;204(2):711–22.

16 Humphreys K, Grankvist A, Leu M, Hall P, Liu J, Ripatti S, et al. The genetic structure of the Swedish population. PLoS One. 2011;6(8):e22547.

17 Kerminen S, Havulinna AS, Hellenthal G, Martin AR, Sarin AP, Perola M, et al. Fine-Scale Genetic Structure in Finland. G3 (Bethesda). 2017;7(10):3459–68.

18 Price AL, Helgason A, Palsson S, Stefansson H, St Clair D, Andreassen OA, et al. The impact of divergence time on the nature of population structure: an example from Iceland. PLoS Genet. 2009;5(6):e1000505.

19 Helgason A, Nicholson G, Stefansson K, Donnelly P. A reassessment of genetic diversity in Icelanders: strong evidence from multiple loci for relative homogeneity caused by genetic drift. Ann Hum Genet. 2003;67(Pt 4):281–97.

20 Van Mol C, de Valk H. Migration and Immigrants in Europe: A Historical and Demographic Perspective. In: Garcés-Mascareñas B, Penninx R, editors. Integration Processes and Policies in Europe: Contexts, Levels and Actors. Cham: Springer International Publishing; 2016. p. 31–55.

21 Tvedt T. Det internasjonale gjennombruddet : fra “ettpartistat” til flerkulturell stat: Dreyers forl.; 2018.

22 Bolstad E. NORSKE POSTNUMMER MED KOORDINATAR: private; 2009 [Collective effort, ~600 anonymous participants]. Available from: https://www.erikbolstad.no/geo/noreg/postnummer/.

23 Purcell S, Neale B, Todd-Brown K, Thomas L, Ferreira MA, Bender D, et al. PLINK: a tool set for whole-genome association and population-based linkage analyses. Am J Hum Genet. 2007;81(3):559–75.

24 Manichaikul A, Mychaleckyj JC, Rich SS, Daly K, Sale M, Chen WM. Robust relationship inference in genome-wide association studies. Bioinformatics. 2010;26(22):2867–73.

25 Alexander DH, Novembre J, Lange K. Fast model-based estimation of ancestry in unrelated individuals. Genome Res. 2009;19(9):1655–64.

26 Lazaridis I, Patterson N, Mittnik A, Renaud G, Mallick S, Kirsanow K, et al. Ancient human genomes suggest three ancestral populations for present-day Europeans. Nature. 2014;513(7518):409–13.

27 Weale ME. Quality control for genome-wide association studies. Methods Mol Biol. 2010;628:341–72.

28 Price AL, Weale ME, Patterson N, Myers SR, Need AC, Shianna KV, et al. Long-range LD can confound genome scans in admixed populations. Am J Hum Genet. 2008;83(1):132–5; author reply 5-9.

29 Reich D, Thangaraj K, Patterson N, Price AL, Singh L. Reconstructing Indian population history. Nature. 2009;461(7263):489–94.

30 Browning BL, Zhou Y, Browning SR. A One-Penny Imputed Genome from Next-Generation Reference Panels. Am J Hum Genet. 2018;103(3):338–48.

31 Browning BL, Browning SR. Improving the accuracy and efficiency of identity-by-descent detection in population data. Genetics. 2013;194(2):459–71.

32 Duforet-Frebourg N, Blum MG. Nonstationary patterns of isolation-by-distance: inferring measures of local genetic differentiation with Bayesian kriging. Evolution. 2014;68(4):1110–23.

33 Browning SR, Browning BL. Accurate Non-parametric Estimation of Recent Effective Population Size from Segments of Identity by Descent. Am J Hum Genet. 2015;97(3):404–18.

34 Waples RS, England PR. Estimating contemporary effective population size on the basis of linkage disequilibrium in the face of migration. Genetics. 2011;189(2):633–44.

35 Wang J, Santiago E, Caballero A. Prediction and estimation of effective population size. Heredity (Edinb). 2016;117(4):193–206.

36 Browning SR, Browning BL. Rapid and accurate haplotype phasing and missing-data inference for whole-genome association studies by use of localized haplotype clustering. Am J Hum Genet. 2007;81(5):1084–97.

37 Tremblay M, Vezina H. New estimates of intergenerational time intervals for the calculation of age and origins of mutations. Am J Hum Genet. 2000;66(2):651–8.

38 Petkova D, Novembre J, Stephens M. Visualizing spatial population structure with estimated effective migration surfaces. Nat Genet. 2016;48(1):94–100.

39 Martin AR, Karczewski KJ, Kerminen S, Kurki MI, Sarin AP, Artomov M, et al. Haplotype Sharing Provides Insights into Fine-Scale Population History and Disease in Finland. Am J Hum Genet. 2018;102(5):760–75.

40 Palamara PF, Lencz T, Darvasi A, Pe’er I. Length distributions of identity by descent reveal fine-scale demographic history. Am J Hum Genet. 2012;91(5):809–22.

41 Xue Y, Mezzavilla M, Haber M, McCarthy S, Chen Y, Narasimhan V, et al. Enrichment of low-frequency functional variants revealed by whole-genome sequencing of multiple isolated European populations. Nat Commun. 2017;8:15927.

42 Lamnidis TC, Majander K, Jeong C, Salmela E, Wessman A, Moiseyev V, et al. Ancient Fennoscandian genomes reveal origin and spread of Siberian ancestry in Europe. Nat Commun. 2018;9(1):5018.

43 Huyghe JR, Fransen E, Hannula S, Van Laer L, Van Eyken E, Maki-Torkko E, et al. A genome-wide analysis of population structure in the Finnish Saami with implications for genetic association studies. Eur J Hum Genet. 2011;19(3):347–52.

44 Njolstad PR, Andreassen OA, Brunak S, Borglum AD, Dillner J, Esko T, et al. Roadmap for a precision-medicine initiative in the Nordic region. Nat Genet. 2019;51(6):924–30.

